# IPF Patient Derived Lung Macrophages are Crucial In Signaling Fibroblast Activation and Disease Pathogenesis

**DOI:** 10.64898/2026.11.18.767401

**Authors:** Abhalaxmi Singh, Preeti Yadav, Nithish Prasad, Anas Mohammad, Sukanta Das, Prerak Gupta, Aritra Bhattacharyya, Mazharul Maishan, Hiroki Taenaka, Jae-Won Shin, Michael A. Matthay, Paul J. Wolters, Ki-Wook Kim, Mallar Bhattacharya, Asrar B. Malik

## Abstract

The role of lung macrophage-generated paracrine signals such as TGFβ1 in activating fibroblasts in the pathogenesis of Idiopathic pulmonary fibrosis (IPF) in humans remains uncertain. Here, employing lung macrophages derived from Idiopathic pulmonary fibrosis (IPF) patients and genomic, proteomic and cell function analysis of macrophages, we showed increased generation of TGFβ1 over a period of 48 hours indicative of aberrantly activated macrophages. In cell culture studies, we demonstrated that macrophages markedly activated lung fibroblasts. By inducing actin reorganization in fibroblasts coupled to increased membrane tension, cell contractility, and expression of α-smooth muscle actin (SMA) and fibroblast activation protein (FAP). Genomic analysis further revealed augmented collagen synthesis, extracellular matrix (ECM) crosslinking, and activation of pro-survival pathways. IPF macrophages exhibited a threefold increase in surface expression of the mannose receptor CD206, which we exploited using mannose-decorated nanoparticles to selectively deliver TGFβ1 siRNA. The nanoparticle mediated TGFβ1siRNA delivery prevented these manifestations of fibroblast activation. Genomic analysis showed that TGFβ1 promoted FOXO-mediated expression of COL1A1 in fibroblasts coupled to decreased expression of Matrix Metalloproteinases MMP2 and MMP11. Together, these findings establish macrophage-derived TGFβ1 as a key driver of fibroblast activation in human IPF and demonstrate the therapeutic potential of macrophage-targeted nanoparticle-mediated gene silencing.

**One sentence summary:** Selective nanoparticle-mediated silencing of macrophage-derived TGFβ1 reprograms pathogenic macrophage–fibroblast crosstalk, restores FOXO-dependent homeostasis in fibroblasts, and prevents fibrotic remodeling in human IPF

## INTRODUCTION

Idiopathic pulmonary fibrosis (IPF) remains a fatal disease characterized by progressive accumulation of extracellular matrix (ECM) proteins such as collagen, disruption of alveolar architecture, and worsening tissue compliance (*1–4*). Current therapies are ineffective, and ultimately require lung transplantation (*5, 6*). Work has demonstrated persistent injury/repair cycles and over time tissue scarring and defective repair pathways as the central disease pathogenic features (*3, 7, 8*). Key cell interactions such between macrophages and fibroblasts and between macrophages and alveolar epithelial cells appear to be important (*1, 7*). However, most of the work to date has been in animal models of lung fibrosis such as the extensive use of bleomycin which do not always simulate complexity of the human disease (*9, 10*). There is for example little understood about the role of cell interactions in human IPF disease pathogenesis (*11*). Macrophages produce a repertoire of fibrogenic cytokines, such as Transforming Growth Factor Beta 1 (TGFβ1) and Platelet-Derived Growth Factor (PDGF), which are touted in disease onset and amplification (*1, 7*). The dominant extant hypothesis derived from animal studies is that activated macrophages drive fibroblast proliferation and ECM accumulation in a pervasive cycle of ECM protein generation and tissue remodeling (*1, 12*). However, evidence supporting this hypothesis is lacking for the human disease. Here, using freshly isolated lung macrophages from a cohort of IPF patients (described Table 1), we observed extensive reprogramming lung macrophages as reflected in 3-fold increases in TGFβ1 production and a shift in phenotype of fibroblasts and generated ECM proteins such as collagen and tissue remodeling. Activation of the transcription factor FOXO in human fibroblasts induced by macrophages from IPF patients, which promotes collagen production and impairs ECM degradation may be central signaling mechanism of the disease.

**Table 1:** Donor profile.

| Condition | Age | Gender | Race | Others |
|---|---|---|---|---|
| Control | 49 | F | Hispanic/Latino | smoked cigarettes |
| Control | 53 | F | White | smoked 1.5 ppd for 40 years |
| Control | 35 | M | White | Smoked cigarettes and marijuana. 1/3 ppd for 20 years |
| Control | 13 | F | African American | medical history of asthma, Cause of death: Head injury |
| Control | 47 | F | American Indian | Intracranial hemorrhage/stroke |
| Control | 50 | F | White | Intracranial hemorrhage/stroke |
| Control | 30 | M | White | smoked fentanyl |
| Control | 18 | M | White | smoked Marijuana, cigarettes and vape |
| IPF | 49 | M | Asian | Never smoker |
| IPF | 44 | M | Hispanic | Never smoker, Probable Familial Fibrosis |
| IPF | 67 | F | White | Never smoker |
| IPF | 66 | F | White | Never smoker, on pirfenidone |
| IPF | 69 | F | White | Never smoker, on Nintedanib |
| IPF | 69 | F | White | Never smoker, on Nintedanib |

## RESULTS

### Profiling patient IPF macrophages and delivery of TGFβ siRNA into macrophages using CD206 expression

Studies were made using human IPF lung macrophages to first determine whether these cells could be selectively targeted to study downstream activation of fibroblasts. Here we employed mannosylated albumin nanoparticles (ANP) as described (*13*) to deliver siRNA nucleotides into macrophages. After obtaining human lung tissue, we isolated lung macrophages as described in **Fig. 1a**. The donor profiles of human lung tissue obtained from IPF and control patients are identified in **Table 1**. FACS analysis of IPF patient derived macrophages showed high expression of CD163 (CD163++) as compared to macrophages from non-IPF control lungs **(Fig. 1b**, **1c)**. This finding reflected the known infiltration of monocyte-derived alveolar macrophages (Mo-AM) in lung fibrosis and consistent with role of Mo-AM in IPF pathology (*1, 7, 13, 14*). Interestingly, CD163++ macrophages also showed augmented expression of CD206 **(Fig. 1b**, **1d)** a known mannose receptor (*15*) as compared to non-IPF resident alveolar macrophages. This finding is consistent with single-cell RNA sequencing data from the Rosas and Kaminsky IPF dataset (*9*) **(Table 2**; **Fig. 1e).** That patient IPF lung macrophages expressed high CD206 was propitious in that it allowed using this antigen to target cells as described herein. We used mannosylated albumin nanoparticles (MANPs), shown to target the mannose receptor (CD206) (*13*), the antigen highly expressed in IPF lung macrophages **(Fig. 2a).** Using flurophore AF647 labeled MANPs, we demonstrated increased MANP internalization by CD163++/CD206++ IPF derived lung macrophages as compared to non-IPF controls **(Fig. 2b).**

**Fig. 1.**
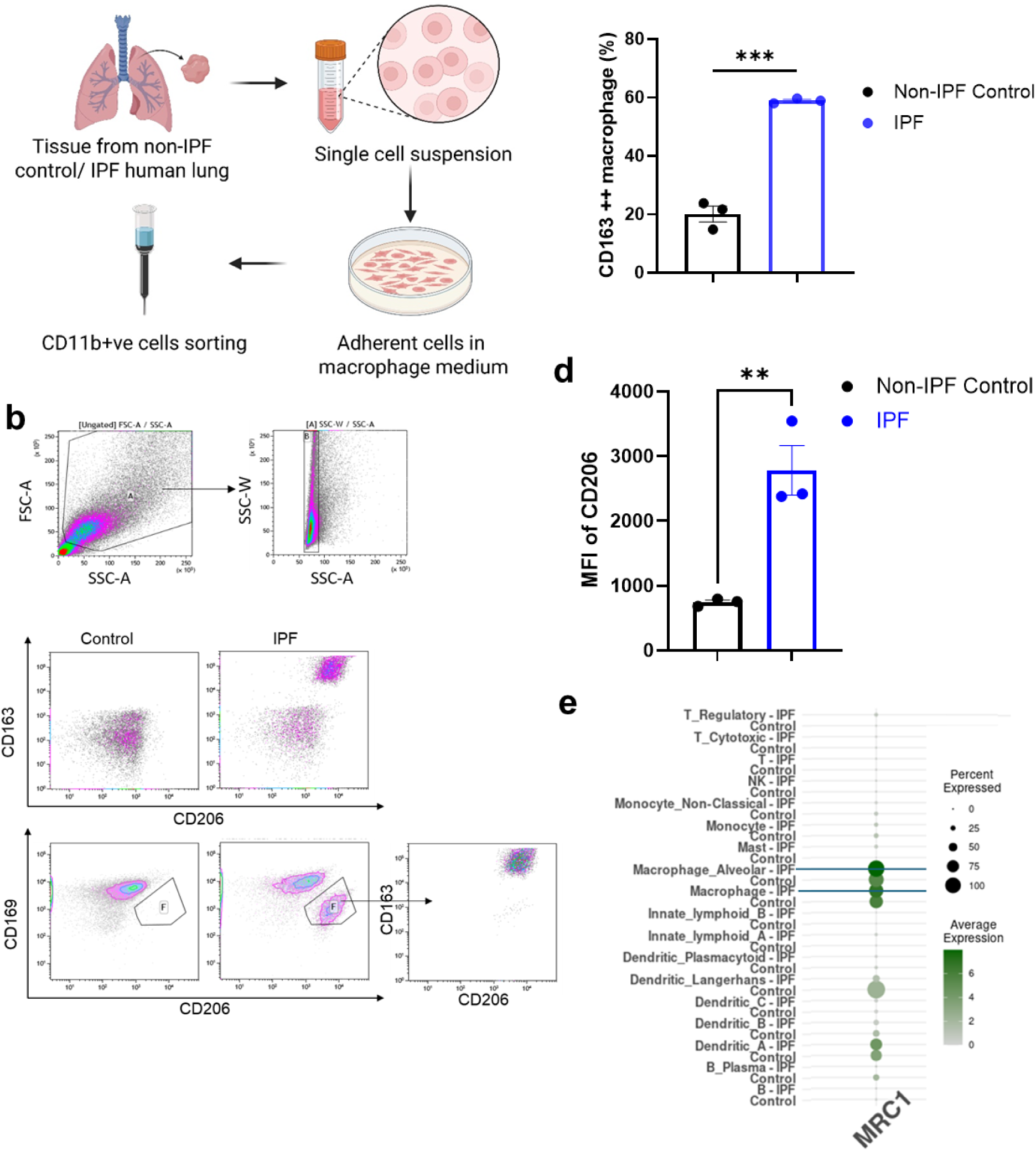
Analysis of macrophages from IPF patient lungs. (a) Schematic for isolation of macrophages from lung tissue. (b) Flow cytometry analysis of macrophages derived from non-IPF control and IPF lungs. I PF lungs showed infiltration of macrophages expressing CD163++ and CD206++. (c) Histoplot showing percentage of infiltrating CD163+ macrophage population in non-IPF control vs. IPF lungs, (d) Histoplot showing mean fluorescence intensity of CD206 in macrophages. (e) Expression of CD206 in alveolar macrophages and lung tissue macrophage population in non-IPF control vs. IPF individuals as obtained from reanalysis of Kaminsky/ Rosas database. Statistical analysis was carried out by t-test. * p<0.05; ** p<0.005 and *** p<0.0005.

**Fig. 2.**
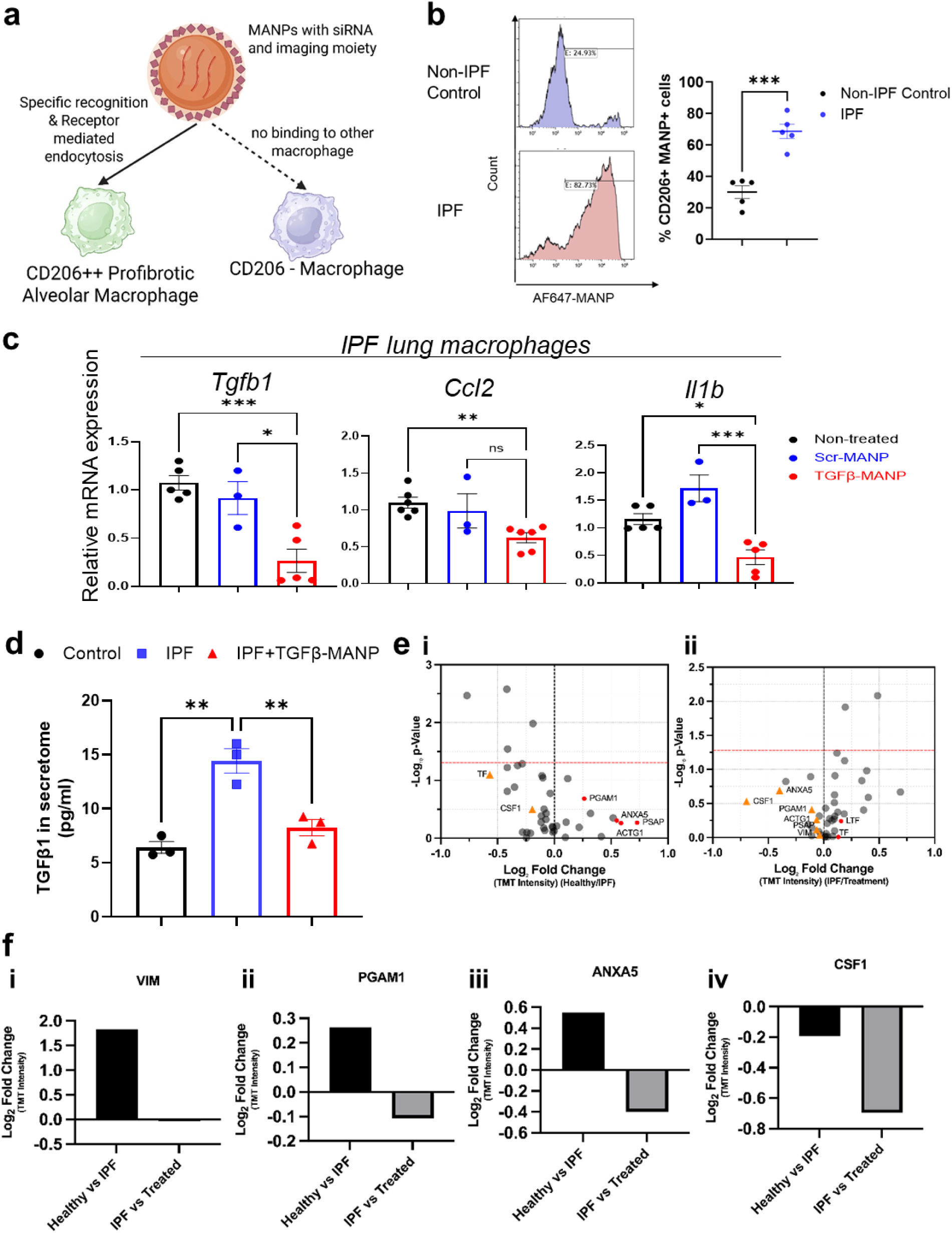
Expression of CD206 is IPF lung macrophages for targeting the pathogenic population. (a) Internalization of mannosylated albumin nanoparticles (MANPS) by non-IPF control and IPF lung macrophages using AF647 fluorophore-labeled MANPs. (b) Table showing fold changes in expression of CD206 (MRC1) and TGFβ1 in macrophages obtained from reanalysis of Rosas/ Kaminsky database. (c) mRNA expression of (i) TGFβ1, (ii) ccl2 and (iii) IL-1β in non-IPF control macrophages, IPF macrophages treated with scrambled siRNA loaded MANPs, and TGFβ1 siRNA loaded MANPs after 48h treatment. Fold change in expression was calculated relative to 18S rRNA gene determined by comparative Ct method. Each data point represents results from macrophages from individual donors. (d) ELISA showing concentrations of TGFβ1 in secretome. (e) Scatter plot analysis showing significant proteins upregulated or downregulated in (i) non-IPF control/ IPF macrophage secretome, (ii) IPF/ TGFβ1 siRNA-loaded MANPs-treated IPF macrophage secretome. (f) Fold changes in expression of key proteins associated with fibrosis progression as identified by proteomic analysis using Scaffold. (i) Vimentin (ii) phosphoglycerate mutase 1 (PGAM1), (iii) Annexin A5 (ANXA5), and (iv) CSF-1. Statistical analysis was carried out using a t-test for two-group analysis and one-way ANOVA for 3-group analysis. * p<0.05; ** p<0.005 and *** p<0.0005.

**Table 2:** Fold change in gene expression by macrophages obtained by reanalysis of Kaminsky-Rosas IPF database.

| ID | HGNC_EnsemblAlt_GeneID | log2FC | log2CPM | PValue | QValue | Contrast | Cluster |
|---|---|---|---|---|---|---|---|
| ENSG00000120708 | TGFBI | 0.56 | 7.52 | 0.00 | 0.00 | IPF/Control | Macrophage |
| ENSG00000120708 | TGFBI | 0.80 | 6.90 | 0.00 | 0.00 | IPF/Control | Macrophage_Alveolar |
| ENSG00000260314 | MRC1 | 0.55 | 9.54 | 0.00 | 0.00 | IPF/Control | Macrophage_Alveolar |

We next used a MANP delivery approach (*13*) to target human IPF lung macrophages. TGFβ siRNA was encapsulated in MANPs to silence TGFβ in human IPF lung macrophages. qPCR showed that TGFβ siRNA-MANPs abrogated expression of TGFβ as well as C-C motif chemokine ligand 2 (ccl2) and interleukin-1 beta (IL-1β) in the IPF macrophages **(Fig. 2c)**. ELISA confirmed reduced TGFβ generation indicating effectiveness of the MANP silencing approach **(Fig. 2d)**.

On analyzing the secretome of TGFβ siRNA-treated lung macrophages, we observed 40 proteins at a cutoff of FDR=5. Several differences were apparent on comparing IPF macrophage vs. TGFβ siRNA-treated IPF macrophage secretomes **(Fig. 2e, f).** Intra-macrophage delivery of TGFβ siRNA into CD163++/CD206++ human IPF macrophages prevented Vimentin (Vim) expression, known to activate inflammatory signaling and differentiate fibroblasts into myofibroblasts (*16*), and induce generation of collagen and remodeling of ECM (*17*) **(Fig. 2f-i)**. TGFβ siRNA also reduced phosphoglycerate kinase1 (PGAM1) expression **(Fig. 2f-ii)** known to occur as a consequence of glycolytic reprogramming of pro-fibrotic macrophages (*18*). In addition, TGFβ siRNA increased Annexin A5 (ANXA5) expression **(Fig. 2f-iii)**, showing that suppressing TGFβ generation prevented macrophage activation (*19, 20*). Colony Stimulating Factor (CSF) expression, linked to macrophage expansion in the fibrotic niche (*21*), was also blocked by suppressing TGFβ generation **(Fig. 2f-iv)**.

### IPF patient derived lung macrophages induce dramatic phenotype shift in fibroblasts

While Mo-AMs are implicated in pathogenesis lung fibrosis (*10, 13*), their role in signaling fibroblast fate shift and ECM deposition is unknown. We thus exposed primary human lung fibroblasts (derived from human lung tissue) (Catalog No. H-6013, Cell Biologics) to IPF lung macrophage secretome **(Fig. 3a)**. Fibroblasts stimulated with IPF macrophage underwent striking morphologic reprogramming characterized by formation of dense actin stress fibers and elongated contractile cell shape changes **(Fig. 3b)**, consistent with transition towards fibrogenic fibroblast phenotype shift (*22, 23*). To address whether TGFβ silencing of IPF macrophages blocked fibroblast phenotype transition, we treated macrophages with TGFβ siRNA-MANPs and applied conditioned media to fibroblasts. Fibroblasts exposed to TGFβ siRNA macrophage secretome showed morphology similar to controls indicating that preventing TGFβ generation blocked aberrant actin remodeling of fibroblasts induced by IPF macrophages **(Fig. 3b)**.

**Fig. 3.**
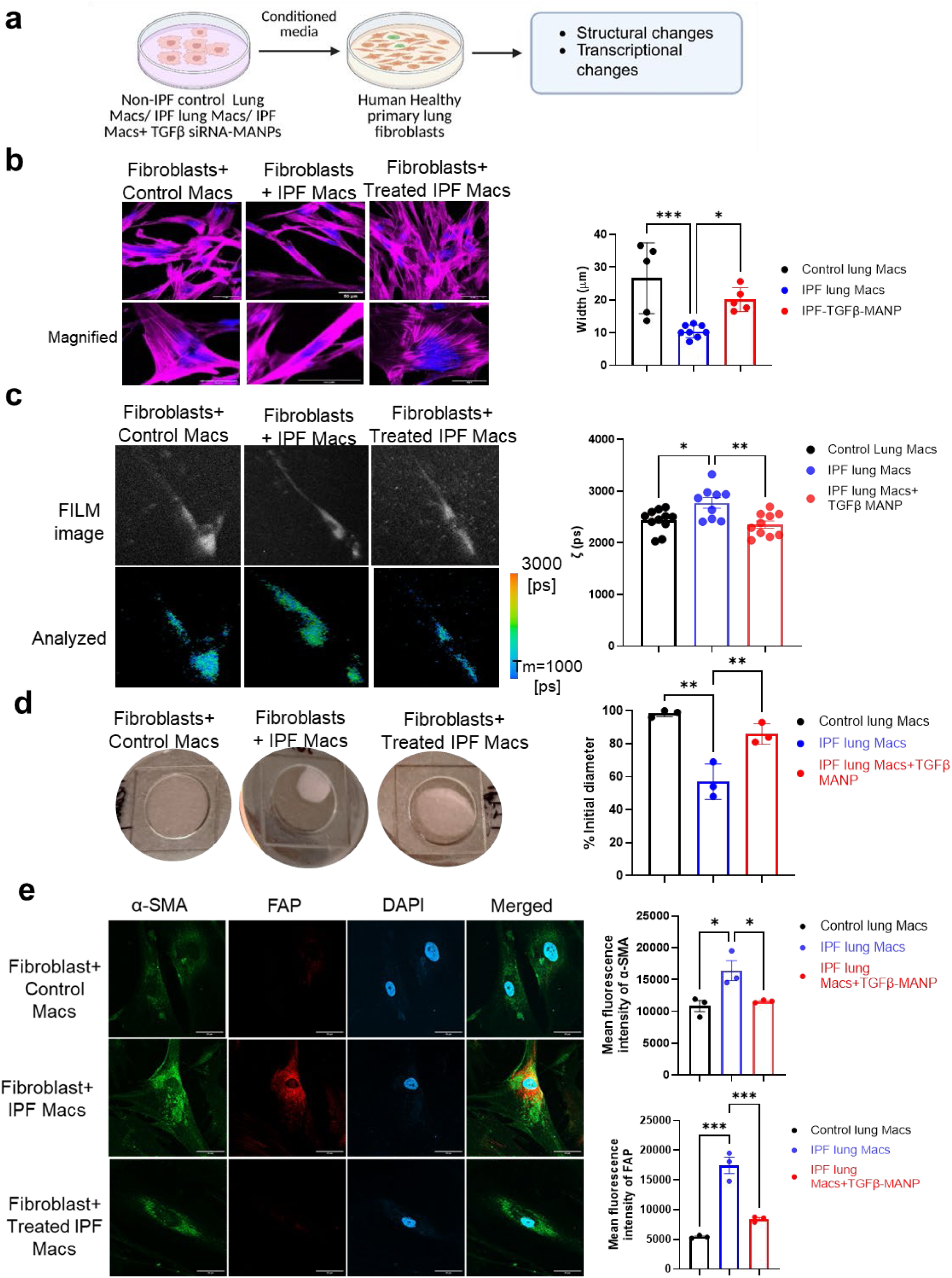
IPF macrophage secretome transfers contractile and tension phenotypes to naïve fibroblasts. (a) Scheme showing experimental setup. Post-seeding ∼20,000 human lung fibroblasts, cells were incubated with control IPF, or TGFβ1 siRNA-loaded MANPs pre-treated with macrophage secretome. After 48h, phenotype of fibroblasts was analyzed. (b) Phalloidin staining was used to determine changes in actin and cell shape of lung fibroblasts incubated with non-IPF control, IPF or TGFβ1 siRNA loaded MANPs pre-treated macrophage secretome. Cell width was measured across the nucleus. (c) Membrane tension was measured using Fluorescence Lipid Tension Reporter that determined alteration in tension landscape of fibroblast membrane. (d) Gel contraction measured contractility of fibroblasts in response to 5h fibrotic macrophage secretome. Enhanced contractility of fibroblasts defined a clear phenotypic shift of fibroblasts towards myofibroblast-like cells. (e) Immunofluorescence staining after 48h treatment with control, IPF or TGFβ1 siRNA loaded MANPs pretreated macrophage secretome. Changes in fibroblasts were quantified after staining with α-SMA and FAP. One-way ANOVA carried out statistical analysis. * p<0.05 and ** p<0.005.

We quantified mechano-structural changes in fibroblasts using FLIM-based membrane tension measurements (*24*). IPF macrophage significantly increased fibroblast membrane tension **(Fig. 3c).** Using the collagen contraction assay, we observed contractility of cells on collagen matrix **(Fig. 3d)**. Both FLIM and contractility assessment of macrophages with TGFβ siRNA-MANPs prevented the shift in fibroblasts to fibrogenic phenotype. Confocal imaging revealed induction of myofibroblast markers alpha-smooth muscle actin (α-SMA) and fibroblast activation protein (FAP) in fibroblasts exposed to IPF macrophage secretome, whereas TGFβ siRNA-prevented upregulation of these markers **(Fig. 3d)**. These findings demonstrate that IPF patient macrophages mediated fibroblast-to-myofibroblast transition secondary to macrophage expression of TGFβ.

### IPF macrophages activate human fibroblast transcriptional program

To determine activation of transcription programs in fibroblasts by IPF macrophages, we performed bulk RNA sequencing of lung fibroblasts treated with macrophage secretome obtained from individual patients. Here 481 genes were significantly altered in the IPF group as compared to control **(Fig. 4a)**. PCA plot showed 2 clusters for the control macrophage secretome vs. IPF macrophage secretome-treated fibroblasts **(Fig. 4b)**. We carried out GO pathway analysis of the datasets. Of the several pathways significantly altered, we focused on those directly linked to fibroblast-mediated ECM modification. IPF macrophages significantly enhanced signaling pathways linked to ECM synthesis, proteoglycan synthesis, and pathways responsible for tissue remodeling. We also observed N-linked glycosylation and proteoglycan synthesis by fibroblasts **(Fig. 4c)** indicative of transition towards myofibroblasts and synthesis of ECM proteins. Some prominently upregulated genes included Type I, II, III collagen, fibronectin 5 and laminin 1 as shown in the Heatmap **(Fig. 4d).** Another important fibroblast modification induced by IPF macrophages was activation of pathways responsible for evasion of cell death programs. Analysis revealed downregulation of pathways linked to apoptosis and upregulation of pathways negatively regulating apoptosis **(Fig. 4c)**. These included upregulation of anti-apoptotic genes BCL2, BCL6, and pro-survival gene BAD and downregulation of apoptotic signal Annexin A1 (ANAXA1) **(Fig. 4d)**. We also observed upregulation of cell signaling pathways TGFβ, JAK-STAT, and IL-17 **(Fig. 4c)**. Interestingly, TGFβ gene as well as cytokine genes, IL-1β and IL-6 were also markedly upregulated **(Fig. 4d)**.

**Fig. 4.**
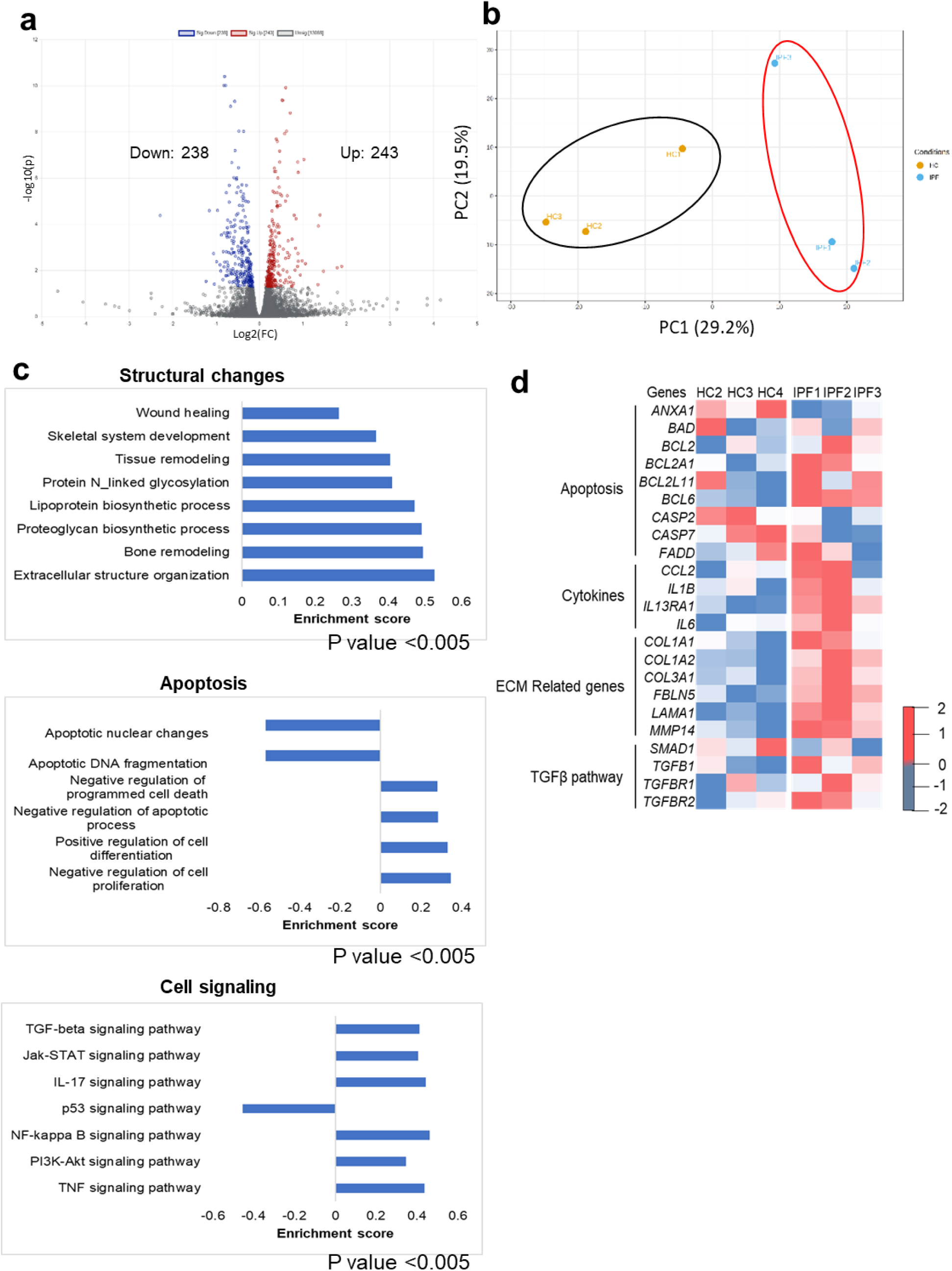
Transcription activation in fibroblasts induced by human IPF derived lung macrophages. (a) Volcano plot showing changes in gene expression in fibroblasts treated with IPF macrophage secretome compared to fibroblasts treated with control macrophage secretome obtained from RNA seq data. X-axis indicates log2 fold change. Red points show upregulated genes and blue points show downregulated genes. (b) PCA plot for RNA-seq data show relationships between samples. (c) Enriched GO terms from up-regulated genes between IPF macrophage secretome vs. control macrophage secretome treated fibroblasts. The enriched pathways were categorized into structural changes, cell death by apoptosis, and signaling pathways which are strongly associated with pro-fibrotic changes occurring in fibroblasts. (d) Heatmap representation of differentially expressed genes, identified by comparing IPF macrophage secretome vs. control macrophage secretome treated fibroblasts. The blue-to-white-to-red gradient represents increased expression of genes with blue representing minimal expression and red representing high expression. Each sample represents lungs from individual donors.

### Specific nanoparticle mediated targeting of patient IPF macrophages abrogates fibroblast phenotype shift

To address transcriptional regulation of IPF macrophage signaling, we analyzed by quantitative PCR expression of pro-fibrotic genes expressed in fibroblasts. We observed increased expression of Actin Alpha 2 (*ACTA2)*, Vimentin (*Vim) and TGFβ1* in IPF secretome-treated fibroblasts **(Fig. 5a)**. These changes were prevented with TGFβ1 siRNA MANP. We compared bulk RNA sequencing data of fibroblasts exposed to control IPF macrophage secretome vs. secretome from TGFβ1 siRNA MANP treated IPF macrophages. We identified 485 genes that significantly altered between two conditions **(Fig. 5b)**. Principal component analysis (PCA) revealed clear segregation of the two groups, indicating distinct transcriptional states driven by TGFβ1 silencing in macrophages as compared to controls **(Fig. 5c)**.

**Fig. 5.**
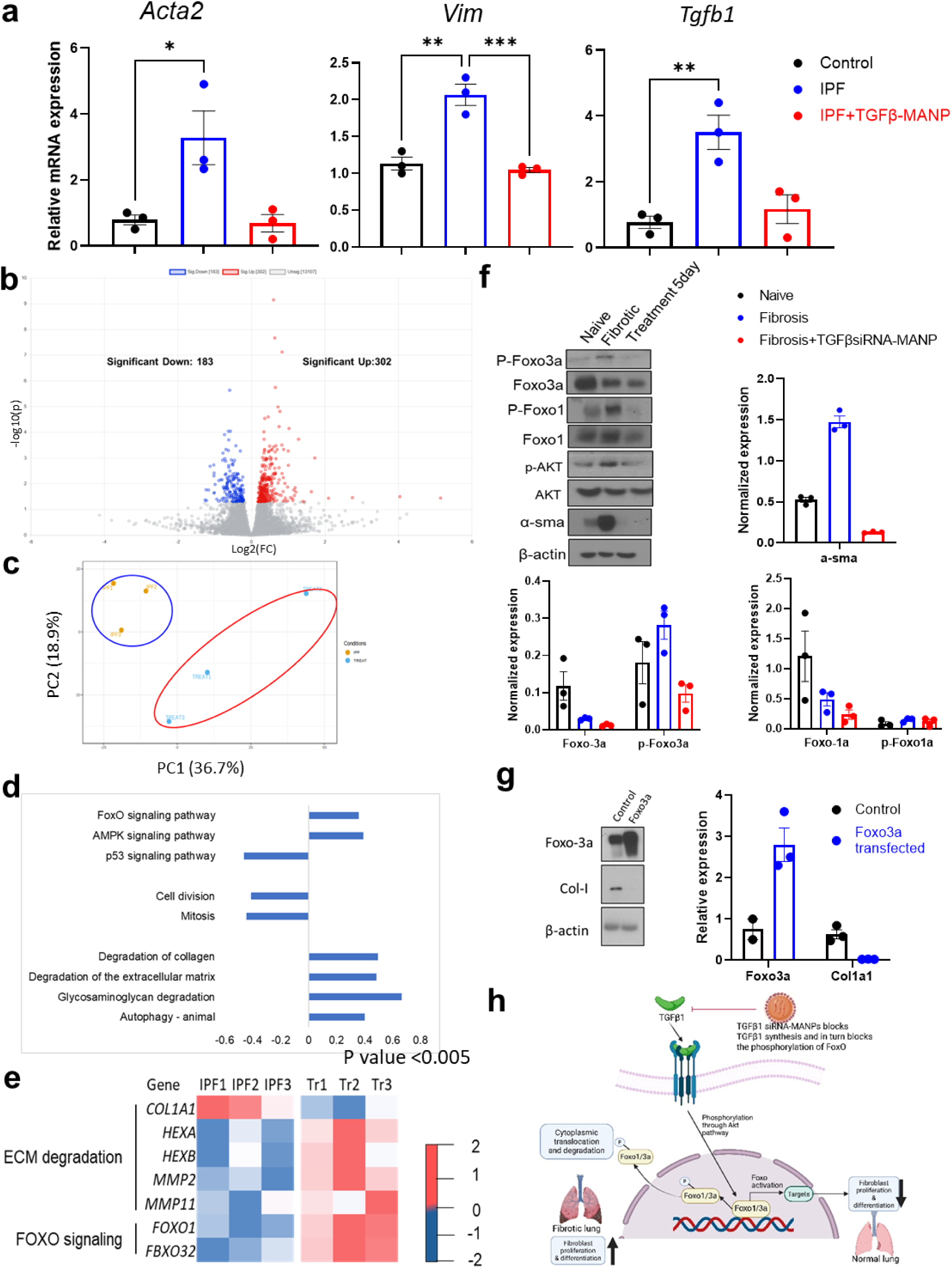
Human IPF lung macrophages loaded with TGFβ1 siRNA-MANP prevent pro-fibrotic transition of fibroblasts. (a) mRNA expression of (i) Acta2, (ii) Vim and (ii) TGFβ1 in control macrophages secretome-treated fibroblasts, IPF macrophage secretome-treated fibroblasts, and TGFβ1 siRNA loaded MANP-treated fibroblasts after 48h period of treatment. Secretomes in each case were collected from macrophages after 48h of culture. Fold change in expression was calculated relative to 18S rRNA gene, as determined by comparative Ct method. Each data point represents results from macrophages from an individual donor. (b) Volcano plot showing overall changes in gene expression in fibroblasts treated with TGFβ1 siRNAloaded MANP-treated IPF macrophage secretome as compared to fibroblasts treated with IPF macrophage secretome, as obtained from RNA seq data. X-axis indicates log2 fold change. Red points indicate upregulated genes, and blue points indicate downregulated genes. (c) PCA plot for RNA-seq data show trelationships between samples. (d) Enriched GO terms from up-regulated genes between TGFβ1 siRNA-loaded MANP-treated IPF macrophage secretome vs. IPF macrophage secretome-treated fibroblasts. The enriched pathways were categorized into structural changes, cell death by apoptosis and signaling pathways which were strongly associated with fibrotic changes in fibroblasts. (e) Heatmap representation of differentially expressed genes, identified by comparing TGFβ1 siRNA loaded MANP-treated IPF macrophage secretome vs. IPF macrophage secretome treated-fibroblasts. The blue-to-white-to-red gradient represents increased expression of genes with blue representing minimal expression and red representing high expression. Each sample represents lungs from individual donors. (f) Western blot analysis of mouse lung fibroblast from naïve, bleomycin-induced fibrotic, and TGFβ1 siRNA-MANP-treated bleomycin challenged-mice. The proteins analyzed included Foxo-3a, p-Foxo3a, Foxo1, p-Foxo1, p-Akt, Akt and α-SMA. β-Actin served as control. The graphs show densitometry analysis of 3 different samples from individual mice. (g) Western blot analysis of IPF lung fibroblast cell line CCL-134 before and after Foxo3a plasmid transfection which shows overexpression of FoxO3a correlates to lower Col1a1 expression (h) Schematic representation of proposed pathway showing TGFβ1 silencing activation of FOXO signaling and in turn shifting balance towards favorable homeostasis. Statistical analysis was carried out using t-test for two-group analysis and one-way ANOVA for 3-group analysis. * p<0.05; ** p<0.005 and *** p<0.0005.

GO analysis highlighted several fibroblast pathways influenced by MANP TGFβ1 siRNA with strongest effects on ECM protein dynamics, collagen turnover, cell–matrix remodeling, and signaling pathways associated with fibrotic progression **(Fig. 5d)**. Notably, fibroblasts in the treatment group showed upregulation of ECM- and collagen-degrading programs, specifically expression of *MMP2* and *MMP11* in parallel with decreased expression of *COL1A1* **(Fig. 5e)**. These signatures are consistent with shift toward matrix breakdown and fibrosis resolution that were aligned with structural and functional reversal of fibroblast activation seen in imaging studies **(Fig. 3)**. Moreover, pathways associated with fibroblast proliferation and profibrotic signaling were broadly downregulated in treatment group, supporting the salutary role of blocking macrophage-driven fibrotic remodeling program by TGFβ1 siRNA MANP. These results together demonstrate that modulating human IPF macrophage activation promotes induction of pro-resolving transcriptional program in fibroblasts and disrupts the central driver of ECM accumulation and fibrogenesis.

### TGFβ1 siRNA targeting of human IPF macrophages prevents FoxO signaling in fibroblasts

RNA-sequencing showed enrichment of FOXO signaling pathway in fibroblasts stimulated with secretome from TGFβ1 siRNA–MANP–treated macrophages as compared with untreated control group **(Fig. 5d).** To investigate transcriptional mechanisms activated by TGFβ silencing of macrophages, we initially used the bleomycin-induced mouse lung fibrosis model (*13, 25*) in which FOXO transcriptional pathway was investigated (*26*). Western blotting demonstrated that fibrotic fibroblasts exhibited high levels of phosphorylated FoxO1a and FoxO3a, modification promoting cytosolic sequestration and proteasomal degradation (*26–28*). In contrast, fibroblasts from TGFβ1 siRNA MANP–treated mice showed inhibition of FoxO1a and FoxO3a phosphorylation **(Fig. 5f)** consistent with activation of FOXO transcriptional program. Furthermore, phosphorylation of FoxO1 and FoxO3a was associated with AKT pathway activation **(Fig. 5f)**. We confirmed similar Foxo3a expression in human fibroblasts. Transfection of FoxO3a plasmid in IPF fibroblasts prevented Col1a expression **(Fig. 5g)**. These findings using IPF patient-derived macrophages further showed that TGFβ silencing prevented fibroblast FOXO phosphorylation, and suppressed fibrotic remodeling **(Fig. 5h)**.

## DISCUSSION

Our results show that specific targeting of a pathogenic population of human lung macrophages using nanoparticle-mediated delivery of TGFβ siRNA prevented human fibroblast activation and deposition of ECM proteins. We observed elevated expression of the mannose receptor CD206 in patient lung macrophages which enabled the specific targeting of these cells (*29*) Furthermore IPF macrophages induced cytoskeleton changes in fibroblasts, reflecting a shift to pro-fibrosis phenotype. This was evident by inhibition of increased membrane tension, contraction of ECM, and elevated expression of α-SMA and FAP in fibroblasts. Detailed transcriptional profiling revealed that fibroblasts shifted genomic profile in response to macrophage release of TGFβ1, resulting in induction of collagen synthetic pathways, ECM crosslinking, activation of proliferative signaling, and metabolic rewiring of fibroblasts consistent with phenotype shift.

A key finding was the observation of increased expression of CD206++ in human IPF patient derived lung macrophages. CD206++ MoAMs have been described in animal studies of lung fibrosis (*10, 13*) and were shown to release pro-inflammatory cytokines and matrix-modifying enzymes (*4, 13, 29, 30*). That CD206 was also highly expressed in IPF patient-derived lung macrophages was fortuitous since it allowed direct targeting of CD206++ cells using mannosylated ANP (MANP) drug delivery probe (*13*). However, it remains unknown whether CD206 itself is a marker of incipient fibrosis or plays a role in disease pathogenesis.

We demonstrated that IPF patient macrophages releases mediators such as TGFβ capable of fibroblast activation and fibrogenesis (*31, 32*). Silencing of TGFβ using siRNA-encapsulated mannosylated albumin nanoparticles reshaped macrophage activation and secretome generation, suppressing both profibrotic and proinflammatory mediator expression, and critically the myofibroblast phenotype shift. The downstream effects of macrophage silencing of fibroblasts included normalization of cytoskeletal architecture, reduced cellular contractility, reduced ECM turnover, and reversal of transcriptional programs favoring ECM degradation and inhibition of fibroblast proliferation.

Another key observation was that restoration of FOXO signaling in fibroblasts downstream of macrophage-dependent TGFβ signaling was crucial in mediating shift in fibroblasts to IPF-inducing myofibroblast phenotype. FOXO1 and FOXO3a upon phosphorylation are degraded, suppressing transcriptional programs that inhibit fibroblast proliferation, oxidative stress, and ECM deposition (*26, 33–35*). Reports from animal studies showed activation of FoxO3 in fibrosis-inducing fibroblasts (*26, 27, 34*) and its crucial role in mediating transition of fibroblasts to myofibroblast phenotype *in vitro* (*26*). We showed that targeted silencing of macrophage-derived TGFβ1 prevented FoxO1 and FoxO3a phosphorylation in human lung fibroblasts from IPF patients, and in thereby restraining fibroblast activation responsible for the disease. Our results using cells from IPF patients support the potential salutatory role of suppressing FOXO activation in fibroblasts as driven by lung macrophages in preventing IPF.

Clinical trials targeting TGFβ however have been equivocal (*32, 36*). The chief reason may be that TGF-β has pleiotropic effects in immune regulation, tissue repair, and epithelial homeostasis (*37*), thus, systemic inhibition may not be beneficial. Another factor may be activation of compensatory signaling networks—such as pathways driven by Platelet-Derived Growth Factor, Connective Tissue Growth Factor, and inflammatory mediators—allowing alternate pro-fibrotic processes to persist (*38, 39*). The present approach of targeting solely lung macrophage population expressing high levels of CD206 is thus more tenable and advantageous.

We took advantage of the albumin nanoparticle-based MANP platform described by us (*13, 40, 41*). This approach was used to identify the upstream role of macrophages in driving IPF in human cells. MANPs provide several advantages such as ease of manufacturing, scalability, and biocompatibility (*13, 40, 42–44*). Albumin is also an established nanocarrier as it shows minimal hepatic accumulation and has fewer hurdles than lipid nanoparticles (*45, 46*). Moreover, mannosylation of albumin nanoparticles ensured preferential uptake by CD206++ macrophages. A possible pitfall of the approach is requirement for high membrane expression of CD206. Fortunately, IPF patient derived lung macrophages showed high expression of CD206 as also previously demonstrated in lungs of IPF patients (*4, 29*). Our results using CD163++/CD206++ macrophages from IPF patients showed generation of TGFβ1 which reprogrammed fibroblasts into highly contractile and ECM-producing cells via activation of FoxO. We showed the potential of targeting CD206++ macrophages using TGFβ1 siRNA loaded macrophages **(Fig. 5h, schematic model)**.

## MATERIALS AND METHODS

### Preparation of mannosylated albumin nanoparticles (MANPs)

Albumin nanoparticles (ANPs) were synthesized as described (*41*) following the desolvation method using Bovine Serum Albumin (BSA MW 66,000Da, Sigma). BSA was diluted in endotoxin-free water (20mg/ml) and reacted with ethanol (3.5ml) with gradual addition over 10m. The formed ANPs were stabilized by adding 38µl glutaraldehyde (25%, Sigma) and stirred overnight. To encapsulate Alexa fluor 647- or Cy5-labeled TGFβ-siRNA was used a uptake identification tracer. The siRNA to be delivered into the macrophages was mixed with 100µl 5% DMSO to which was added BSA solution for 1h. ANPs were collected by centrifugation (15,000g, 20min, 4°C) and washed 3 times by resuspension in endotoxin free water (1mL). After the third wash, the pellet was resuspended in PBS for further use in studies.

Mannosylated albumin nanoparticles (MANP) were then prepared using our published protocol (*13*). Briefly, D-mannose (Sigma) (0.8 M) was dissolved in sodium acetate buffer (pH 4.5) and heated at 60 ℃ for 1h to open the mannose ring and expose the aldehyde group for further conjugation. Then, the cooled down activated mannose was added to ANP suspension (1:5 v/v ratio) in endotoxin free water and left for overnight stirring. Next day MANPs were collected by centrifugation (15,000 g, 20min, 4°C.) and washed 3 times by resuspension in endotoxin free water (1mL). After third wash, the pellet was resuspended in PBS.

### Human lung macrophages and uptake of nanoparticles

All tissue were obtained following IRB protocols and institutional guidelines. Macrophages were isolated from healthy lungs or IPF lung tissue obtained after transplantation or diseased donors. The tissues were digested with collagenase to obtained single cells. The cell suspension was allowed to adhere to culture dishes to eliminate any non-adherent monocytes, neutrophils etc. Only the adherent cells were further processed through CD11b sorting column to enrich the lung macrophage population. The macrophages post identification were kept frozen until use. Lung macrophages from healthy and IPF individuals were revived and cultured for 5 days in 20 % FBS, DMEM media prior to the studies. The lung macrophages were identified by their surface receptors CD14 and CD64 (Biolegend).

The human lung macrophages upon revival were further characterized to identify alveolar macrophages (AMs) or monocyte derived alveolar macrophages (Mo-AMs) by flow cytometry. The AMs are specifically identified by their expression of CD169 (Biolegend0 and Mo-AMs were identified by their expression of CD163 (Biolegend). Further, the expression of CD206 was analyzed in the different lung macrophage subpopulation.

To study uptake of MANP, we AF647-conjugated MANP (1µg/ml MANPs) and incubated these with healthy or IPF lung macrophages for 15 min, which were then washed with PBS and cells were collected by gentle scraping. The cells were then collected centrifugation and analyzed by flow cytometry. For treatment of IPF macrophages. scrambled siRNA or TGFβ si-MANPs (1µg/ml MANPs) were treated to IPF macrophages. To collect the secretome of healthy or IPF or treated IPF macrophages, the conditioned media (secretome) was collected after 48h of culture and filtered through 0.2µm filter and stored at -80℃ for further analysis.

All the flow cytometry analyses were carried out in Flow Fortessa (BD Biosciences) and data were analyzed by Kaluza Analysis 2.1 (Beckman). For analysis, the cells were first gated for live single cells, followed by CD14/ CD64 and then further analyzed for different expressions.

### Macrophage-fibroblasts axis assessment

To evaluate macrophage-fibroblast axis, secretome collected from the healthy/ IPF lung macrophages were added to human lung primary fibroblasts. All the effects were evaluated after 48h of incubation.

### RNA isolation and real-time qPCR

Total RNA was isolated from the macrophage or fibroblast using the RNeasy Plus kit (Qaigen). After quantification by Nanodrop 1000 (Thermo Fisher), 500 ng of RNA were reverse-transcribed into complementary DNA using High-Capacity cDNA Reverse Transcription Kit (Thermo Fischer scientific) according to manufacturer’s instructions. The cDNA obtained was mixed with FastStart Universal SYBR Green Master (Rox) (Milipore Sigma) with specific qPCR primers for qPCR on an ABI Prism 7000 system and analyzed with for relative quantification on the ViiA-7 and Quant studio Real-Time PCR System (Thermo Fisher).

### Bulk RNA-sequencing

Agilent Bio-analyzer was used to check RNA quality and quantity. All samples showed RNA integrity number > 8. RNA-Seq libraries were prepared using Illumina mRNA TruSeq kits according to the protocol by Illumina. Library quality and quantity were checked and the pool of libraries was sequenced using an Illumina HiSeq4000 and Illumina reagents (GSE231219).

### RNA-sequencing

Raw reads were aligned to reference genome mm10 using the STAR (Dobin et al., 2013). ENSEMBL genes were quantified using Feature Counts (Liao et al., 2014). Differential expression statistics (fold-change and p-value) were computed using edgeR (McCarthy et al., 2012) (Robinson et al., 2010), using the generalized linear model (GLM) capability to correct for a batch effect between groups of replicates. P-values were adjusted for multiple testing using the false discovery rate (FDR) correction of Benjamini and Hochberg. Significant genes were determined based on an FDR threshold of 5% (0.05). The data were compared first between healthy vs IPF macrophage secretome treated fibroblasts to know the IPF macrophage effect on fibroblasts. Then, the treatment effect was studies by comparing the IPF macrophage vs TGFβsiRNA-MANP treated IPF macrophage secretome treated fibroblasts. The up- and down-regulated genes from were obtained. Gene ontology biological process (GO BP) pathways were obtained from Express Analyst, and enrichment statistics for up- and down-regulated gene lists were computed using Fisher’s Exact test.

### Macrophage secretome analysis

Sample (5 ug of protein) preparation was carried out with Filter Aided Sample Preparation (FASP) protocol using 10 K NMWL centrifugal filter units. Dithiothreitol was added at a final concentration of 20mM and reduced for 30 minutes. Proteins were then alkylated with IAA at a final concentration of 50 mM IAA for 20 min in the dark. The alkylated proteins were washed with 200uL 8M Urea in 0.1M Ammonium Bicarbonate (ABC) 3 times and equilibrated with 0.1M ABC 3 times prior to tryptic digestion. Proteins were digested using Trypsin at an enzyme to protein ratio of 1:20 in 50uL 0.1M ABC buffer at 37 °C for overnight. The digested peptides were collected by centrifugation at 14,000*g* for 20 min followed by two elution with 50μL of 50mM ABC and one elution with 50uL of 0.5M NaCl. The eluted peptides were desalted using Oasis PRiME HLB (30mg) and dried. The samples were labeled with TMT10plex Isobaric Label Reagent Set (ThermoFisher, Waltham, MA) followed by the manufacturer’s instructions. The scheme for TMT labeling of samples is shown in Table 2. Briefly, 5 ug samples were re-suspended in 50uL 50mM Triethylammonium bicarbonate (TEAB) buffer, and 400ug TMT reagents (Channel 1 to Channel 9) in anhydrous acetonitrile were added into each sample. After 2 hours incubation at room temperature, the reaction was quenched with 4uL of hydroxylamine for 15 min. All 9 samples were combined, dried and desalted with a primed HLB 96 well plate for the high-pH reverse phase (HPRP) fractionation. 45µg pooled TMT peptides were fractionated by off-line high-pH reverse phase (HPRP) liquid chromatography. The peptides were separated into 80 fractionations with XBridge BEH C18 Column, 130Å, 3.5μm, 4.6mm X 250 mm (Waters). Every 11th fraction was concatenated together and 10 concentration fractions were dried. Then, peptides were resuspended in 60 µL of 5% acetonitrile, 0.1% formic acid buffer for LC-MS analysis.

For data analysis, spectra were searched against Uniprot human database using Mascot daemon (2.6.0) with the following parameters; parent mass tolerance of 10 ppm, constant modification on cysteine alkylation, variable modification on methionine oxidation, and deamidation of asparagine and glutamine. Search results were entered into Scaffold Q+S software (v5.0.0, Proteome Software, Portland, OR) for compilation, normalization, and comparison of spectral counts, etc. TMT purity correction was applied with the reporter ion isotopic distributions. The filtering criteria of protein identification were a 1% false discovery rate (FDR) of protein and peptide with 1 minimum peptide count.

### Immunofluorescence and confocal microscopy

After 48h of macrophage secretome exposure, culture lung fibroblasts were washed with PBS followed by 10 min Fc block. Then surface staining was done with Fibroblast activation protein (FAP) primary antibody for 30min. The cells were then fixed with 4% paraformaldehyde (Sigma #P6148) and permeabilized with 0.05% Triton X-100 (Fisher Scientific #BP151–100). After a wash with PBS/Tween-20 (Fisher Scientific #BP337–100), cells were blocked with 5% BSA (Sigma) in PBS/Tw-20 for 1 h at room temperature. Afterward, cells were incubated with anti-alpha-SMA primary antibodies and incubated overnight at 4 °C. The next day, slides were washed and incubated with the fluorescence-conjugated secondary antibody. Images were taken with a confocal microscope LSM880 (Zeiss) and analyzed by ImageJ. For Phalloidin staining to visualize the actin filaments and structure of the fibroblast, the staining was done after the fixation step.

### Gel contraction study

Collagen gel contraction was carried out as following previously published protcol by Lee et al.(*47*), with minor modifications Briefly, Type I collagen, rat tail was diluted in macrophage-conditioned media (healthy, IPF, or IPF macrophages treated with TGFβ-siRNA-loaded MANPs) to a final concentration of 2 mg/mL. Primary Lung Fibroblasts were trypsinized and resuspended at 2 × 10⁵ cells/mL. Equal volumes of the collagen solution and cell suspension were mixed to yield a final concentration of 1 mg/mL collagen containing 1 × 10⁵ cells/mL per well. The gels were allowed to polymerize at 37°C, and gel diameters were measured over 48 hours. Contraction was calculated relative to the initial gel diameter. All experiments were performed in triplicate and independently repeated three times for each condition.

### Secretome ELISA

The level of TGFβ in the secretome from the healthy, IPF or treated IPF macrophages was quantified by ELISA kit (Thermo) according to the manufacturer’s instructions.

### Western blotting

Cells were washed with cold PBS and lysed in RIPA buffer containing a protease inhibitor cocktail. After clearing the lysate, samples were prepared in sample loading buffer containing SDS and 2-mercaptoethanol. Proteins were separated on a 10% gel and transferred to PVDF membranes. Membranes were blocked in either 5% skimmed milk or 3% BSA (for phosphorylated protein detection) and incubated with primary antibodies. The primary antibodies anti-P-FoxO3a, anti-FoxO3a, anti-P-Foxo1, anti-Foxo1, anti-sma and anti-actin. Following washing, membranes were incubated with appropriate HRP conjugated secondary antibodies and proteins were visualized using Thermo Pico PLUS substrate.”

### Statistical analysis

Statistical analyses were performed using Graph Pad Prism, version 9.0 and with corrections for multiple comparisons whenever appropriate. Data are displayed as mean ± SEM. Statistical significance was evaluated as indicated in the figure legends and is represented by the following scheme: ∗ p≤0.05, ∗∗ p≤0.01, ∗∗∗ p≤0.001, ∗∗∗∗ p≤0.0001.

## Acknowledgements

The authors thank Donor West for supplying the human lungs from brain dead donors for these studies. We thank Fluorescence Imaging Core for confocal microscopy imaging and analysis and Flow Cytometry Core for cell sorting and flow cytometry data acquisition. We thank Bisheng Zhou for helping during RNA sequencing sample preparation and Mark Maienschein-Cline and Jason Michael Wood for the initial analysis of RNA sequencing data. Bioinformatics analysis in the project described was performed by the UIC Research Informatics Core, supported in part by NCATS through Grant UL1TR002003. The secretome analysis was performed by the Mass Spectrometry Core in the Research Resources Center of the University of Illinois at Chicago. The authors would also like to acknowledge the NIH s10 shared instrumentation grant (1S10OD027016-01) for supporting this work. The studies were supported by NIH grants P01HL151327-05. M.B. was supported by institutional/discretionary funds from the University of California, San Francisco.

## Author Contributions

A.S. and A.B.M. conceived the project. A.S., J.W.S., M.B. K.K. and A.B.M. designed the experiments and evaluated the data. A.S., P.Y., N.P., A.M., S.D., P.G. performed the experiments and analyzed the data. A.B., M.M., H.T., M.M., P.W. M.B. received the human lung samples and provided the macrophages for the experiment. A.S., M.B., K.K. and A.B.M. wrote the initial manuscript draft. All authors reviewed the manuscript and provided critical feedback.

